# Phylogenomic resolution of sea spider diversification through integration of multiple data classes

**DOI:** 10.1101/2020.01.31.929612

**Authors:** Jesús A. Ballesteros, Emily V.W. Setton, Carlos E. Santibáñez López, Claudia P. Arango, Georg Brenneis, Saskia Brix, Esperanza Cano-Sánchez, Merai Dandouch, Geoffrey F. Dilly, Marc P. Eleaume, Guilherme Gainett, Cyril Gallut, Sean McAtee, Lauren McIntyre, Amy L. Moran, Randy Moran, Pablo J. López-González, Gerhard Scholtz, Clay Williamson, H. Arthur Woods, Ward C. Wheeler, Prashant P. Sharma

**Affiliations:** Department of Integrative Biology, University of Wisconsin–Madison, Madison, WI, USA; Queensland Museum, Biodiversity Program, Brisbane, Australia; Zoologisches Institut und Museum, Cytologie und Evolutionsbiologie, Universität Greifswald, Greifswald, Germany; Senckenberg am Meer, German Centre for Marine Biodiversity Research (DZMB), c/o Biocenter Grindel (CeNak), Martin-Luther-King-Platz 3, Hamburg, Germany; Biodiversidad y Ecología Acuática, Departamento de Zoología, Facultad de Biología, Universidad de Sevilla, Sevilla, Spain; Department of Biology, California State University-Channel Islands, Camarillo, CA, USA; Départment Milieux et Peuplements Aquatiques, Muséum national d’Histoire naturelle, Paris, France; Institut de Systématique, Évolution, Biodiversité (ISYEB), Sorbonne Université, CNRS, Concarneau, France; Department of Biology, University of Hawai’I at Mānoa, Honolulu, HI, USA; Institut für Biologie, Vergleichende Zoologie, Humboldt-Universität zu Berlin, Berlin, Germany; Division of Biological Sciences, University of Montana, Missoula, MT, USA; Division of Invertebrate Zoology, American Museum of Natural History, New York City, NY, USA

**Keywords:** arthropods, Pycnogonida, mitogenome, ultraconserved, diversification

## Abstract

Despite significant advances in invertebrate phylogenomics over the past decade, the higher-level phylogeny of Pycnogonida (sea spiders) remains elusive. Due to the inaccessibility of some small-bodied lineages, few phylogenetic studies have sampled all sea spider families. Previous efforts based on a handful of genes have yielded unstable tree topologies. Here, we inferred the relationships of 89 sea spider species using targeted capture of the mitochondrial genome, 56 conserved exons, 101 ultraconserved elements, and three nuclear ribosomal genes. We inferred molecular divergence times by integrating morphological data for fossil species to calibrate 15 nodes in the arthropod tree of life. This integration of data classes resolved the basal topology of sea spiders with high support. The enigmatic family Austrodecidae was resolved as the sister group to the remaining Pycnogonida and the small-bodied family Rhynchothoracidae as the sister group of the robust-bodied family Pycnogonidae. Molecular divergence time estimation recovered a basal divergence of crown group sea spiders in the Ordovician. Comparison of diversification dynamics with other marine invertebrate taxa that originated in the Paleozoic suggests that sea spiders and some crustacean groups exhibit resilience to mass extinction episodes, relative to mollusk and echinoderm lineages.

## Introduction

Pycnogonida (sea spiders), the sister group to the remaining Chelicerata, are exclusively marine arthropods ranging from one to 750 mm in size (figure 1). The body architecture of sea spiders is unusual, with a typically very small body that is dwarfed by the much longer legs (hence, the alternate name “Pantopoda”, or “all legs”), into which diverticula of major organ systems emanate. Sea spiders are found throughout the world’s oceans from the intertidal zone to abyssal depths, but are especially abundant and diverse in polar benthic communities. In contrast to many invertebrate groups that flourish in the tropics, the peak of sea spider diversity is concentrated in the Southern Ocean, which also harbors multiple cases of gigantism in distantly related species [1,2].

**Figure 1.**
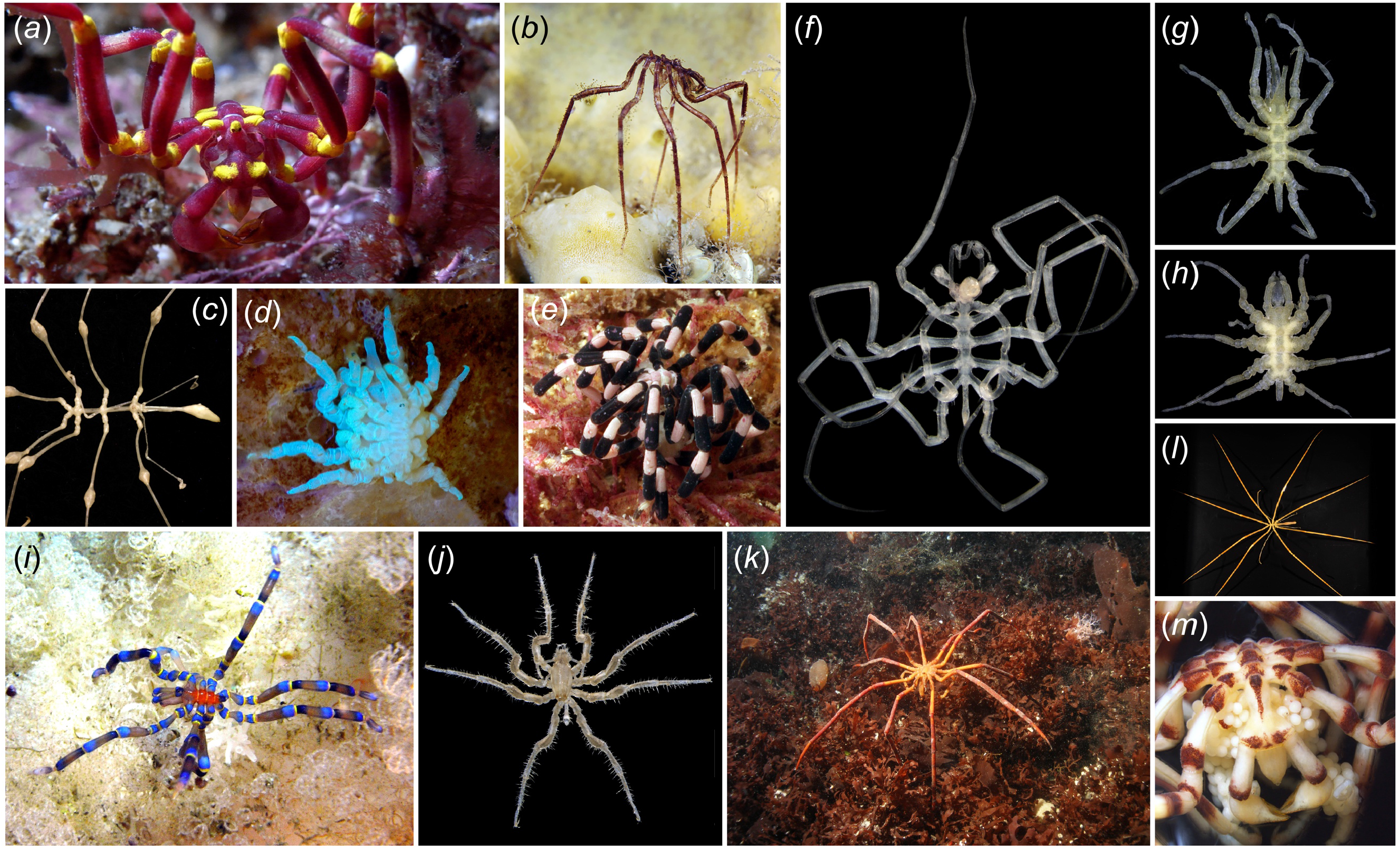
Exemplars of sea spider diversity. (*a*) *Meridionale harrisi* (Callipallenidae). (*b*) *Nymphon grossipes* (Nymphonidae). (*c*) *Rhopalorhynchus magdalenae* (Colossendeidae). (*d*) Copulating pair of *Pycnogonum litorale* (Pycnogonidae) with UV illumination. (*e*) *Stylopallene* sp. (Callipallenidae), photograph by Iain Gray. (*f*) *Nymphonella tapetis* (Ascorhynchidae *sensu lato*). (*g*) *Austrodecus glaciale* (Austrodecidae). (*h*) *Rhynchothorax australis* (Rhynchothoracidae). (*i*) *Anoplodactylus evansi* (Phoxichilidiidae). (*j*) *Cilunculus armatus* (Ammotheidae), (*k*) *Decolopoda australis* (Colossendeidae), photograph by Andrei Utevsky. (*l*) *Colossendeis megalonyx* (Colossendeidae). (*m*) Male of *Meridionale* sp. (Callipallenidae) with egg clutch.

Pycnogonids typically have four pairs of walking legs attached to the small body; the cephalon bears an anterior triradiate proboscis, and three pairs of cephalic appendages, the chelifores, palps, and ovigers. Extant Pycnogonida lack both a segmented opisthosoma (abdomen or posterior tagma) and thus, the segmentally iterated opisthosomal respiratory organs that are found in other chelicerates; sea spiders respire instead via cuticular gas exchange, with peristaltic contractions of the gut facilitating oxygen transport through the body [3,4]. Ovigers, a type of modified leg unique to Pycnogonida, are used for grooming and by the males to carry egg masses (figure 1*m*) [1]. A remarkable exception from the conserved body architecture are genera with supernumerary body segments (resulting in 10-legged species), which occur in three families (Colossendeidae, Pycnogonidae, and Nymphonidae), and one genus of colossendeids is even characterized by 12 legs [5]. Beyond this, the cephalic appendages show generally a high degree of variation. Families are often distinguishable by the number of articles in the palps and ovigers [6], and in several cases, adults may lack one or more cephalic appendage types altogether.

Early conceptions of sea spider phylogeny envisioned a gradual reductive trend characterized by unidirectional, stepwise losses of appendage types [7–9]. Phylogenetic investigations of sea spider relationships based on anatomical data [6] or combined analyses of morphology and molecular sequence data [5] suggested instead that reduction of appendages occurred independently across the phylogeny, but tree topologies were highly discordant between data partitions. Subsequent approaches to infer sea spider relationships under model-based approaches [10–12] were repeatedly frustrated by the instability of basal relationships, which are attributable to two possible causes.

First, efforts to infer the phylogeny of Pycnogonida have been based on a small number of loci (one to six genes; figure 2). These datasets consisted largely of genes that evolve at high rates (e.g., mitochondrial genes cytochrome *c* oxidase subunit I and 16S rRNA) or those that evolve at uninformatively low rates (e.g., nuclear ribosomal genes) [5,10–12]. Separately, mitochondrial genes of sea spiders exhibit well-known lineage-specific compositional biases [11]. Thus, datasets based on fast-evolving mitochondrial genes have exhibited limited utility in resolving Paleozoic relationships of various invertebrate groups, and the placement of sea spiders within Chelicerata specifically [13,14].

**Figure 2.**
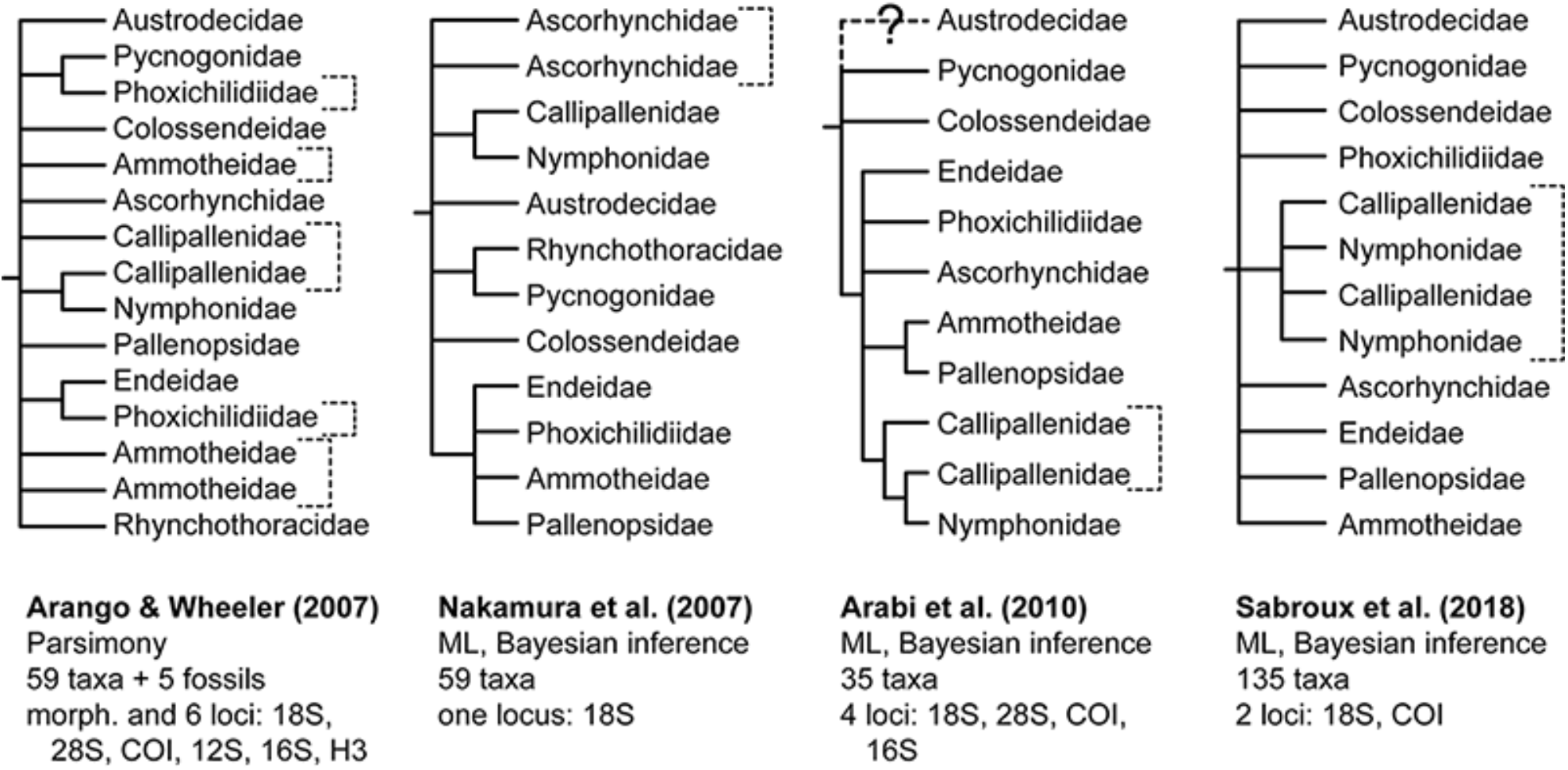
Historical hypotheses of higher-level sea spider relationships based on molecular sequence data. Nodes lacking support (<50% bootstrap; <95% posterior probability) or conflicting between analyses in each study have been collapsed. Brackets correspond to non-monophyletic lineages.

Second, previous phylogenetic studies have omitted or poorly sampled two small-bodied families of sea spiders, Austrodecidae and Rhynchothoracidae [11,12]. Austrodecidae (figure 1*g*; approximately 60 species in two genera) are distinguished from other sea spiders by the annulation of the proboscis. Little is known about their biology, as most austrodecids are small-bodied species (< 5 mm) that are infrequently encountered [1,2]. Even less understood are species of Rhynchothoracidae (figure *1*h; approximately 20 species in one genus), which typically do not exceed one millimeter in length [1,6].

The lack of a robust sea spider phylogeny has hindered inferences of major macroevolutionary trends in the group, such as latitudinal biogeographic patterns, larval developmental mode, and the evolution of appendages, body plans, and neuroanatomical structures [15–17]. While phylotranscriptomic approaches have proven remarkably effective for resolving relationships of numerous chelicerate groups [18,19], the inaccessibility of rare sea spider lineages has obviated RNA-Seq-based approaches, as cryptic and small-bodied species are often not identified in ethanol-preserved samples until weeks to years after their initial collection. Moreover, sea spider specimens are often covered with epibionts, and the sea spider digestive system extends into all but the two most distal podomeres of the legs. As a result, RNA-Seq-based approaches carry high risks of contaminations from epibionts and gut contents, especially for small-bodied species.

Resolving sea spider relationships thus requires an approach that (1) is suited for specimens of various ages in museum collections, (2) overcomes the limitations of mitogenomes and Sanger-sequenced loci, (3) amplifies sea spider sequence specifically, and (4) is robustly applicable to all families of sea spiders, including small-bodied lineages. To surmount these challenges, we undertook a target capture sequencing approach toward generating a robustly resolved phylogenetic backbone for Pycnogonida. We present here the first phylogenomic tree of sea spiders sampling all extant families. To place this branch of the Tree of Life in a temporal context, we inferred for the first time the age of the crown group Pycnogonida using a node dating approach under a Bayesian inference framework.

## Materials and Methods

A list of taxa sampled from field expeditions, museum collections, and multiple deep sea cruises is provided as electronic supplementary material, table S1. Taxonomic sampling consisted of 89 sea spiders; outgroups consisted of 14 Arachnida (including one Xiphosura, which has recently been shown to be nested within the arachnids [20, 21]), three Myriapoda, three Pancrustacea, and one Onychophora. Methods for molecular work, library assembly, multiple sequence alignment, trimming, and phylogenetic inference are detailed in the electronic supplementary material, text S1. Briefly, we designed probes for mitochondrial genomes and nuclear exons using genomic resources previously generated by us and/or mitogenomes in GenBank. For UCEs, we deployed probes from the UCE Arachnida 1.1Kv1 bait set [22]. Probe synthesis, automated library preparation, and paired-end sequencing (2 × 150 bp) was performed on the Illumina Hi-Seq 2500 platform through RAPiD Genomics (Gainesville, FL, US). We included the nuclear ribosomal genes 5.8S rRNA, 18S rRNA, and 28S rRNA, which were obtained as bycatch. Outgroup taxa were subsequently added into the alignments using available genomic resources.

Gene trees were inferred using IQ-TREE v.1.6 [23] with the best-fitting model selected by ModelFinder (-m MFP) [24]. Four matrices were constructed for our main analyses. Matrices 1-3 were constructed using taxon occupancy thresholds of 50%, 33%, and 25%, respectively. Matrix 4 consisted of loci stipulated to sample at least one Austrodecidae (the putative sister group to the remaining sea spiders). Final alignments were analyzed under maximum likelihood using IQ-TREE v.1.6 with best-fitting models per locus. Due to ongoing debate over the benefits of model-fitting, we additionally analyzed our datasets using a unique GTR + Γ4 model for each locus as well. Nodal support was estimated using bootstrap resampling frequency with 1000 ultrafast bootstrap replicates in IQ-TREE. Approximately Unbiased (AU) tests of monophyly were performed using in-built tools in IQ-TREE for selected phylogenetic hypotheses using Matrices 1 and 4. Bayesian inference analyses were performed using PhyloBayes-mpi v.1.7 with a CAT + GTR + Γ4 model [25].

Phylogenomic estimation of divergence times was estimated using a node dating approach with MCMCTree [26] on two datasets (Matrices 1 and 3), implementing a likelihood approximation of branch lengths using a multivariate normal distribution [27]. Fossils used to inform the dating consisted of 11 outgroup and four ingroup node calibrations. A list of these calibrations and their use, as well as an overview of all sea spider fossils evaluated for this purpose, is provided in electronic supplementary material, text S2. Both the independent rates and correlated rates clock models were used to infer node ages with Matrices 1 and 3, under the maximum likelihood tree topology inferred for Matrix 3 with a unique substitution model per locus (figure 3). Analyses of diversification rates through time were performed using Bayesian Analysis of Macroevolutionary Mixtures (BAMM v.2.5.0) [28], as specified in in electronic supplementary material, text S1.

**Figure 3.**
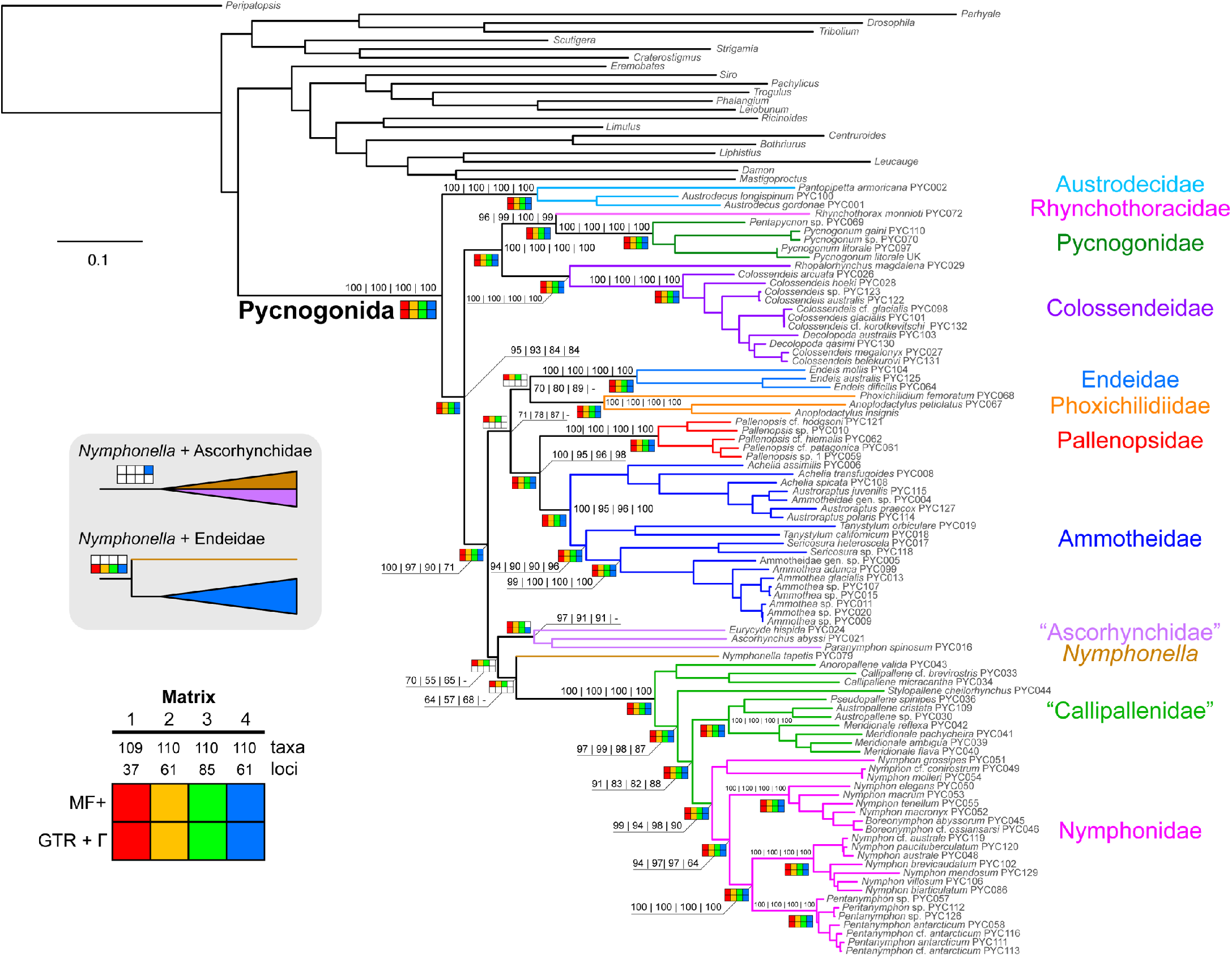
Phylogenomic relationships of Pycnogonida based on maximum likelihood analysis of Matrix 3 (ln*L* = −475920.91). Colors of branches correspond to families (right). Numbers on nodes indicate bootstrap resampling frequencies for Matrices 1-4 with model-fitting using ModelFinder. Bottom left: Sensitivity plot indicating design of matrices and phylogenomic analyses. Inset (gray background): Alternative placements of *Nymphonella tapetis.*

Micro-computed tomography scans were performed with an Xradia MicroXCT-200 (Carl Zeiss Microscopy). Detailed procedures and scan settings are provided in electronic supplementary material, text S1. Processing and 3D visualization of image stacks was performed with Imaris v. 7.0.0. (Bitplane AG, Switzerland), as described previously [29].

## Results

### (a) Phylogenomic relationships of Pycnogonida

Maximum likelihood analyses of Matrices 1-4 consistently recovered the monophyly of Pycnogonida with maximal nodal support (bootstrap resampling frequency [BS]) (figure 3). Austrodecidae was recovered in all analyses as the sister group to the remaining sea spiders with support (BS = 84-95%). At the next internal node, we recovered a clade comprised of Colossendeidae, Pycnogonidae, and Rhynchothoracidae (BS = 100%), with unambiguous support for the sister group relationship of the latter two families (BS = 96-100%). All other families were robustly recovered as a distal clade (BS = 74-100%).

As shown in figure 3, ML analyses of Matrices 1-4 with model-fitting using ModelFinder recovered Endeidae and Phoxichilidiidae as sister taxa (BS = 70-89), the latter family encompassing its two constituent genera *Phoxichilidium* and *Anoplodactylus*, which had been challenged in a previous molecular analysis [5]. Endeidae + Phoxichilidiidae are in turn sister group (BS = 71-87%) to a clade comprised of the mutually monophyletic Pallenopsidae and Ammotheidae (BS = 95-100%). The remaining sea spider lineages consisted of a monophyletic Nymphonidae unambiguously nested within Callipallenidae (BS = 97-99%); and this lineage in turn subtended with weak support by a grade comprised of *Nymphonella tapetis* (BS = 57-68%) and a paraphyletic Ascorhynchidae *sensu stricto* (BS = 55-70%). Ascorhynchidae consistently included the putative ammotheid genus *Paranymphon* (BS = 91-97%). Comparable results were recovered in PhyloBayes-mpi analyses, with support (electronic supplementary material, figure S1).

Model-fitting strategy did not affect the backbone of this tree, save for the placement of *Nymphonella.* Assigning a unique GTR + Γ model to each locus for Matrices 1-4 consistently recovered *Nymphonella* as the sister group of Endeidae. Matrix 4, which was constructed on the basis of the inclusion of autrodecid sequence in all loci (rather than predicated on dataset completeness), alternatively recovered *Nymphonella* as nested within Ascorhynchidae when the partition models were developed using ModelFinder (figure 3). The PhyloBayes topology for Matrix 3 recovered the same backbone tree, but with *Nymphonella* as the sister group to the clade formed by Endeidae, Phoxichilidiidae, Ammotheidae, Pallenopsidae, Callipallenidae, and Nymphonidae (without support; electronic supplementary material, figure S1).

Barring the placement of *Nymphonella* and its attendant topological instability, all analyses congruently resolved the monophyly of all families, excepting Rhynchothoracidae (one terminal only), Ammotheidae (due to the placement of *Paranymphon)*, and Callipallenidae (consistently recovered as paraphyletic with respect to Nymphonidae). To assess support for non-monophyly of Callipallenidae, we performed AU tests, using a constraint tree that enforced the mutual monophyly of Callipallenidae and Nymphonidae. The monophyly of Callipallenidae was rejected under both Matrix 1 (Δln*L* = 101.92, *p* = 1.67 × 10^−3^) and Matrix 4 (Δln*L* = 132.34, *p* = 3.27 × 10^−4^).

Most genera sampled with two or more taxa were also recovered as monophyletic. Exceptions included *Decolopoda* (a result attributable to the lack of any loci that include both *Decolopoda* species) and *Achelia* (sampled with three terminals). Our results corroborated the non-monophyly of *Colossendeis* due to the nested placement of species with supernumerary segments (e.g., the genera *Decolopoda* and *Dodecolopoda*) [30], and additionally reveal here the non-monophyly of *Nymphon* due to the nested placement of *Boreonymphon* and the 10-legged genus *Pentanymphon.*

### (b) Comparative assessment of phylogenetic data classes

Performance measures for phylogenetic data classes consisted of number of taxa sampled, alignment length, GC content, Robinson-Foulds distance (RF), weighted Robinson-Foulds distance (wRF), and evolutionary rate (electronic supplementary material, text S1). A detailed dissection of dataset performance is provided in electronic supplementary material, text S3. Briefly, the mean number of taxa captured per locus was highest for the mitochondrial and nuclear ribosomal genes (68.2 and 85.7, respectively) and comparable for targeted exons, albeit with high variance (mean=50.9, σ^2^=20.9) (electronic supplementary material, figure S2). UCE loci bore the most missing data, with an average of 22.2 terminals per locus. Assessment of congruence between gene trees and the pruned species trees showed that targeted exons and UCEs had comparable distributions of RF and wRF distances. Evolutionary rates of targeted exons and UCEs were comparable to each other, and intermediate between mitochondrial genes and nuclear ribosomal genes (electronic supplementary material, figure S3).

### (c) Age and tempo of sea spider diversification

Phylogenomic estimation of divergence times under a correlated rates clock model and the most complete matrix (Matrix 1) recovered a Late Cambrian to Ordovician age for the crown group of Pycnogonida (median: 481 Mya; 95% HPD interval: 451-504 Mya) (figure 4*a*). Comparable ages were recovered for the same dataset under an uncorrelated rates clock model (median: 464 Mya; 95% HPD interval: 427-500 Mya). Diversification of sea spider families was estimated to have occurred during the Paleozoic under both clock models. BAMM analysis of sea spider diversification revealed no evidence of rate shifts within Pycnogonida, recovering instead a monotonic and near-constant process of diversification (electronic supplementary material, figure S5).

**Figure 4.**
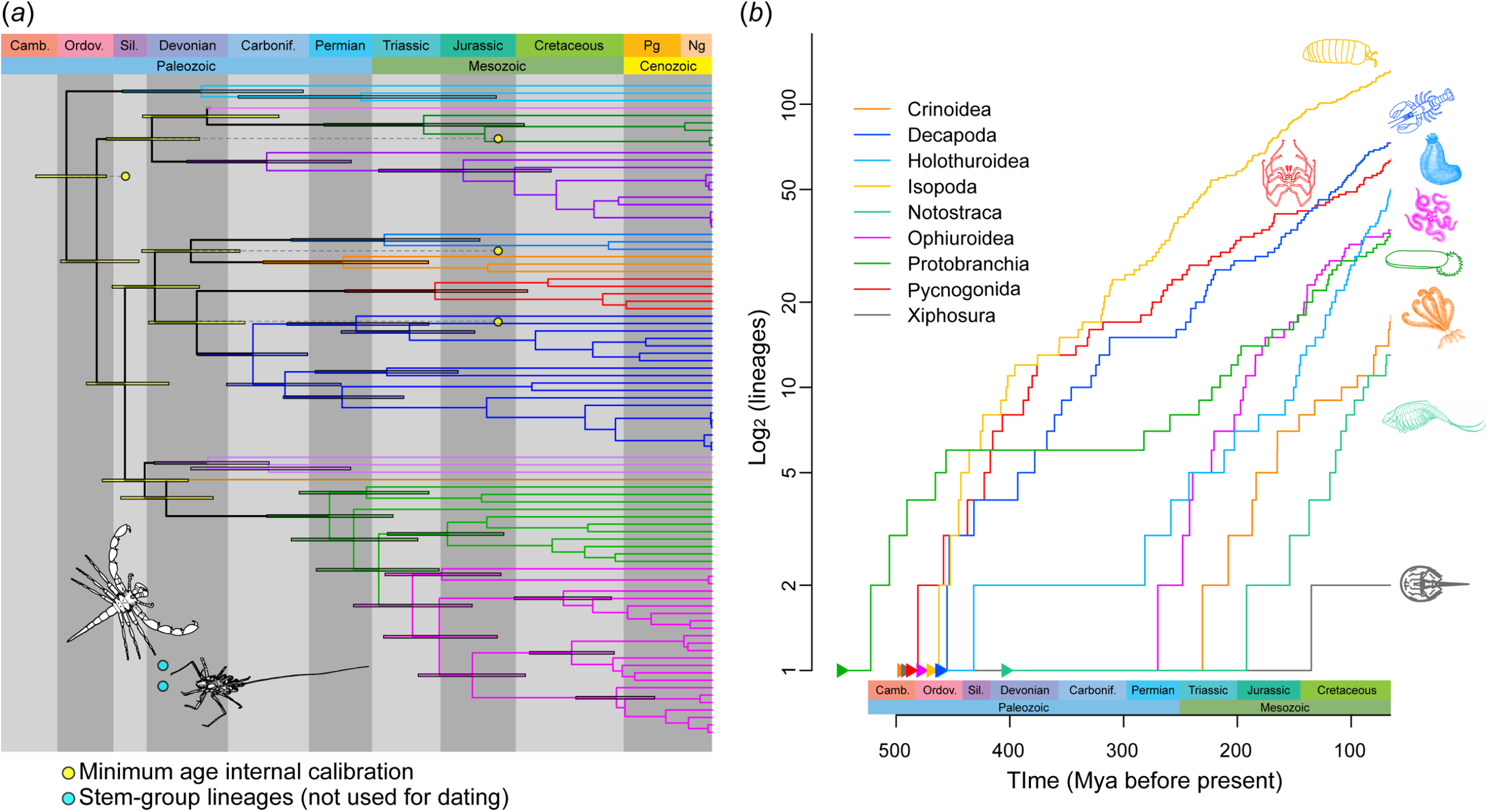
(*a*) Phylogenomic dating of sea spiders based on the most complete data matrix and a correlated rates molecular clock model. Colors of branches and 95% HPD intervals correspond to families, as in figure 3. Line drawings (inset) correspond to stem-group fossils *Palaeoisopus problematicus* (top) and *Flagellopantopus blocki* (bottom). (*b*) Log lineage through time trajectories for selected Paleozoic aquatic taxa (sources in text). Branching times are truncated at the Cenozoic to mitigate under-sampling of recent diversity and/or oversampling of intraspecific terminals.

## Discussion

### (a) A phylogenomic view of higher-level sea spider relationships

Traditional views of sea spider evolution suggested that the sister group of the remaining sea spiders was Nymphonidae, based on its generalized appendage morphology) [7], or possibly Ammotheidae (previously thought to include *Eurycyde*) [31] based on anatomical similarities between ammotheids and fossils like *Palaeoisopus problematicus.* The species tree that we produced decisively recovered the enigmatic family Austrodecidae as the sister group to the remaining sea spiders, consistent with an earlier proposal to separate austrodecids from all other Pycnogonida as the sole member of the suborder Stiripasterida [8,32]. The recovery of a sister group relationship of Pycnogonidae and Rhynchothoracidae is consistent with the morphology of these families, as well as a previous analysis of 18S rRNA [9,32]. Overall, our dataset established a stable backbone topology for the remaining families (figure 3). Only the position of *Nymphonella tapetis* engendered topological discordance across the datasets.

Establishing the sequence of basal branching families facilitates reconstruction of evolutionary transformation series for major character systems. Upon reconstructing adult appendage characters on our tree topology, we established that adult chelifores have been lost in a grade of families at the base of extant Pycnogonida (figure 5). Given some ambiguity in the state of the chelifore in some fossil taxa as well as uncertainty about their phylogenetic placement, our results suggest that the presence of an adult chelifore may not be the unambiguous ancestral condition for crown-group sea spiders. The sister group relationship of Endeidae and Phoxichilidiidae accords with the shared loss of palps and female ovigers in these families. We also consistently obtained the non-monophyly of Callipallenidae, recapitulating a result that has been suggested, albeit with weak support, in previous analyses of one to six loci [5,10,11]. In contrast to previous topologies, we recovered the monophyly of Nymphonidae as a derived lineage within the callipallenids across all datasets and analyses. This result calls for systematic revision of these two families.

**Figure 5.**
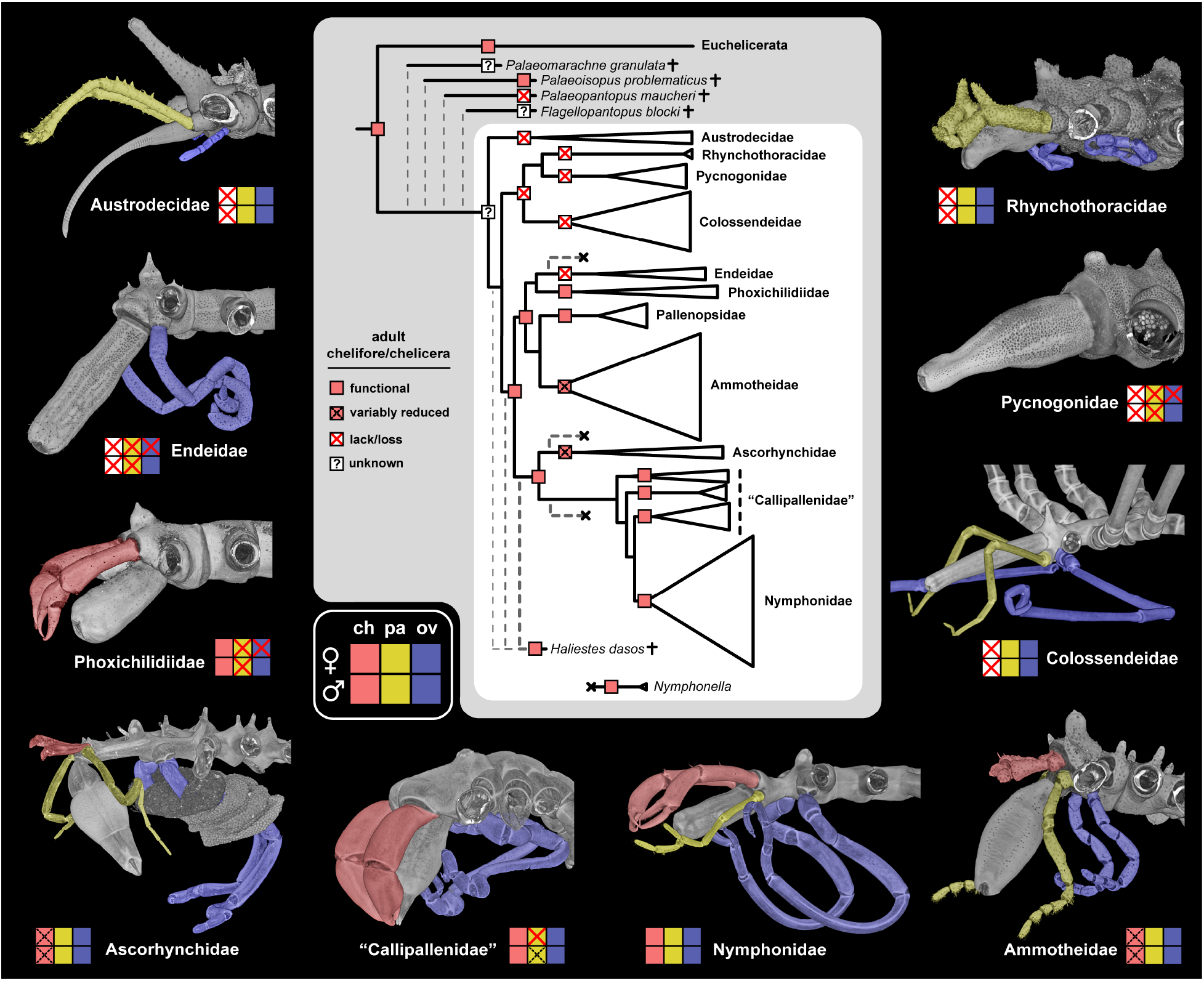
Cephalic appendage evolution in sea spiders, with emphasis on the chelifore. Reconstruction of adult chelifores is mapped on the topology obtained herein, with addition of fossil stem group and crown group representatives. Ancestral state reconstruction is based on equally weighted parsimony. Note the omission of functional adult chelifores in colossendeids with supernumerary segments (a derived state within the genus *Colossendeis*). Specimens in counter-clockwise order from top left: male *Austrodecusglaciale*, male *Endeis spinosa*, female *Phoxichilidium femoratum*, egg-bearing male *Ascorhynchus ramipes*, male *Stylopallene cheilorhynchus*, male *Nymphon gracile*, female *Ammothella longipes*, female *Colossendeis angusta*, female *Pycnogonum litorale*, female *Rhynchothorax australis.*

While genera were largely recovered as monophyletic, those defined to accommodate the condition of supernumerary segments (e.g., *Decolopoda, Pentanymphon*) were nested within larger genera of eight-legged species (e.g., *Colossendeis, Nymphon*). The exception was *Pentapycnon*, although our sampling of the family Pycnogonidae remains too sparse to test the monophyly of *Pycnogonum* (ca. 100 described species). Lineages with supernumerary segments are remarkable in an evolutionary context; in other Chelicerata, segment number tends to be fixed within a given extant order, as exemplified by such groups as scorpions and harvestmen. Changes in the segmentation of the prosoma (the appendage-bearing tagma of chelicerates) are especially rare (e.g., the synziphosurines *Offacolus kingi* and *Weinbergina opitzi* [33]), yet all sea spiders are distinguished from the remaining Chelicerata by the presence of an additional oviger-bearing segment (the post-tritocerebral segment, which bears a walking leg in all other chelicerates). The mechanistic basis of supernumerary segment addition is currently an enigma, due to limited developmental genetic resources for sea spiders [34,35]. Future work should prioritize the gap segmentation genes of sea spiders, especially those functionally linked to gap segmentation phenotypes in chelicerate model species [36–38].

### (b) *Nymphonella* and the composition of Ascorhynchidae

Surprisingly, the position of the putative ascorhynchid *Nymphonella tapetis* was unstable to analytical treatment of the dataset, a result that cannot be attributed to missing data alone *(N. tapetis* was represented in 47-59% of loci across Matrices 1-4). *Nymphonella* is clearly distinguished from other sea spiders by anterolaterally projecting chelifores, annulated distal podomeres of the first walking leg, and a bulbous proboscis (figure 1*f*). Previous efforts to infer the placement of *Nymphonella* have recovered limited support for its placement or for the monophyly of Ascorhynchidae, though *Nymphonella* has only been represented by 18S rRNA in such studies [12,39]. *N. tapetis* may therefore constitute a rogue taxon, an inference corroborated by its variable placement across data partitions generated in this study (electronic supplementary material, figure S4).

By contrast, we recovered support for a nested placement of the putative ammotheid genus *Paranymphon* [32] within Ascorhynchidae *sensu stricto* (represented by the genera *Eurycyde* and *Ascorhynchus*). *Paranymphon* has never been previously sequenced, but like *Nymphonella*, this terminal exhibited topological incongruence across data classes, being recovered within ammotheids by the mitochondrial data partition (albeit being represented therein by only three mitochondrial genes), and with ascorhynchids by UCEs and exons (13 and 17 loci, respectively).

Given the uncertainty surrounding the composition of Ascorhynchidae, future systematic efforts should target deeper sequencing of the genera *Nymphonella* (three described species) and *Paranymphon* (six described species).

### (c) Combining data classes overcomes limitations of individual partitions

Historical efforts to infer higher-level relationships of sea spiders have leaned heavily on the Sanger-sequenced nuclear ribosomal and mitochondrial markers, but these have not yielded a stable sea spider phylogeny to date [5,11,12]. Ideally, new markers for filling this gap should exhibit evolutionary rates intermediate between slow-evolving nuclear ribosomal genes and fast-evolving mitochondrial markers. Targeted exons and UCEs sequenced in this study exhibit rates that propitiously fall exactly within this desired range (electronic supplementary material, figure S3).

Problematically for the UCE dataset, capture efficiency of UCE probes was much lower than other data partitions (electronic supplementary material, figure S2). This phenomenon appears to be partly linked to the availability of DNA in a given extraction; a previous study on spider systematics that deployed the same UCE probe set discovered low recovery rates for extractions with small quantities of DNA [40]. Low quantity of DNA was unavoidable for such minute and rare lineages as *Rhynchothorax* and austrodecids. However, coverage inefficiency of the arachnid UCE probe set was systemic for sea spiders, even when ample DNA was available for large-bodied species. As a result, of an initial 230 UCEs amplified, we discarded 56% of alignments where fewer than six sea spider terminals were obtained. The remaining 101 UCE loci bore high proportions of missing data, in contrast to the targeted exons. Similar outcomes have been reported for palpimanoid spiders, a group that had the advantage of a well-annotated theridiid spider genome for validation of probe design [40].

Nevertheless, we observed high informativeness of the UCE loci despite missing data, as inferred from distributions of RF and weighted RF distances. The concatenated UCE tree, while incongruent with the species tree, recovered such higher-level groupings as Nymphonidae nested within Callipallenidae, and Colossendeidae sister group to Rhynchothoracidae + Pycnogonidae. Given the promise of this phylogenetic data class, efforts to improve the recovery of UCE datasets in sea spiders should target the generation of high-quality sea spider genomes, with downstream improvements in the design of sea spider-specific UCE probes. Such strategies have been shown to overcome limitations inherent to the arachnid UCE bait set for spiders [41].

### (d) The first molecular dating of Pycnogonida reveals ancient diversification and monotonic evolutionary rates

Generally, fossil records for invertebrates (especially marine arthropods) are scarce. The appearance of a Cambrian fossil resembling a sea spider early developmental instar, together with the exquisite preservation of the fossil *Haliestes dasos*, clearly point to an ancient origin of sea spiders before the Silurian (electronic supplementary material, text S2). However, a fossil record dating to the Paleozoic is not dispositive of ancient diversification of the crown group. Evolutionary relicts like Xiphosura (horseshoe crabs) appeared early in the fossil record (oldest fossil belonging to the Ordovician), but survived to the present as only four species that diverged in the Cretaceous [42]. Molecular divergence time estimation for such groups as Crinoidea (feather stars) and Ophiuroidea (brittle stars) have shown that both these echinoderm groups diversified in the wake of the end-Permian mass extinction [43,44], having survived this extinction episode as a single lineage that subsequently recovered some fraction of its diversity (a “revenant” taxon *sensu* [45]). By contrast, marine invertebrate groups like Holothuroidea (sea cucumbers) [46] and Protobranchia (protobranch bivalves) [47] diversified in the Paleozoic but retain the signature of the end-Permian mass extinction in their dated phylogenies, which manifests as a low diversification rate (the plateau of an anti-sigmoidal curve in a log-lineage through time plot) until the beginning of the Mesozoic (figure 4*b*) [47].

To date, molecular divergence time estimation has never been performed for sea spiders. It was previously postulated that sea spiders diversified relatively recently (in the Mesozoic), but this speculation was based solely on the branch length (i.e., substitutions per site) subtending Pycnogonida in a molecular phylogeny, rather than a parametric molecular dating approach or the use of fossil calibrations [11]. Our divergence time estimation unambiguously recovered an Ordovician age of sea spider diversification, a result that is independent of both the deployment of *Haliestes dasos* as a calibration prior, and the choice of clock model. Ancient diversification of Pycnogonida during the Ordovician is consistent with their fossil record (e.g., Jurassic sea spiders that are assignable to families; electronic supplementary material, text S2), and further suggests that Devonian sea spiders with opisthosomal segments (e.g., *Flagellopantopus, Palaeoisopus*) constitute stem lineages that diverged from extant Pycnogonida before the Ordovician and thereafter went extinct. A parallel evolutionary history has been reconstructed for spiders and their extinct sister group Uraraneida, which was recently shown to have survived at least until the Cretaceous [48].

Comparison of lineage accumulation through time for selected marine invertebrate groups reveals the marked difference between the evolutionary history of sea spiders and other Paleozoic fauna (figure 4*b*) [42–44,46,47,49,50]. In contrast to groups like Protobranchia and Crinoidea, Pycnogonida exhibited a static diversification regime, with a monotonic process of slowing diversification rate since initial divergence, and no evidence of rate shifts, by comparison to other Paleozoic taxa (electronic supplementary material, figure S5). The lack of any signature of major mass extinction events in the early (pre-Mesozoic) evolutionary history of extant sea spider and crustacean lineages is unexpected. In contrast to groups like bivalves and echinoderms, sea spiders do not form large calcified hard parts (valves or tests) whose deposition was severely affected by cataclysmic historical environmental changes (e.g., during the end-Permian event) [51]. However, this differing composition of hard parts does not account for the Paleozoic origin and post-Permian diversification of aquatic arthropod groups like Xiphosura and Notostraca [42, 52]. Moreover, the cuticle of decapods is indeed biomineralized, with denser cuticle associated with higher calcification in benthic Decapoda [53]. Apropos, the log-lineage through time trajectory for Decapoda does indeed exhibit a small decline in lineage accumulation rate immediately preceding the Triassic (figure 4*b*), but this result was only observed under divergence time estimation under one of two clock models [49].

We therefore postulate that the resilience of sea spiders is not exclusively due to their lack of a calcified exoskeleton, but could be also attributable to higher diversification rates in cooler regions, as has previously been shown in marine arthropod groups like Anomura [54,55]. In deep-sea isopods, diversification in the deep sea occurred in parallel with anoxic events, with the earliest radiation dated to the early Permian and subsequent episodes of rapid colonization and radiation [50]. Such processes could partly explain the oddities of sea spider biogeography (such as the concentration of their diversity in Antarctica), but the role that the Southern Ocean has played as a potential center of endemism is not clear based on our results. In the present study, which is focused on higher-level phylogenetic relationships, we lack sufficiently broad geographic sampling to assess the biogeography of sea spiders and a putative role for the polar regions as the driver of sea spider diversity in lower latitudes. Population genetic works have begun tackling such questions at shallower taxonomic scales (within species or species groups), with high levels of precision [30,56–62].

We add here the caveat that analyses of diversification rates are highly contingent on sampling intensity. Therefore, intensive taxonomic sampling, in tandem with phylogenomic approaches and the establishment of high-quality genomic resources for sea spiders, will be essential vehicles toward unlocking diversification dynamics and evolutionary history of Pycnogonida.

## Ethics

Specimens from multiple deep sea cruises were collected under the auspices of the United States Antarctic Program, in compliance with the US Antarctic Conservation Act. All diplomatic formulars for sampling the Icelandic economic zone were given for cruises M85/3 (IceAGE1) and POS456 (IceAGE2). Specimens from Australia were collected under permit no. 15111 of the Tasmanian Department of Primary Industries, Parks, Water and Environment and Marine Parks permit G08/27858.1.

## Data accessibility

Sequence data: GenBank. Multiple sequence alignments, partition files, phylogenetic trees, probe sequences: Dryad Digital Repository.

## Author contributions

C.P.A, W.C.W., and P.P.S. conceived of the study. Specimens were collected and/or contributed by C.P.A., G.B., S.B., E.C.S, G.F.D., M.P.E., G.C., L.M., A.L.M. R.M., G.G., H.A.W., P.L.G., and P.P.S. A new mitogenome of *P. litorale* was contributed by G.S. for bait design. Coordination of sample sorting from the DIVA3 and IceAGE cruises was performed at the DZMB in Hamburg by S.B. Taxonomic identification from IceAGE and DIVA3 collections was performed by C.P.A, G.B., E.V.W.S., and P.P.S. E.V.W.S. and P.P.S performed molecular work. E.V.W.S. performed imaging of specimens used for extraction. G.B. performed μCT scans and their subsequent processing and visualization. J.A.B., C.E.S.L., M.D., S.M., J.S., C.W., and P.P.S performed bioinformatic and phylogenetic analyses. W.C.W. and P.P.S funded the work. P.P.S drafted the manuscript, and all authors edited and approved the final content.

## Competing interests

We declare we have no competing interests.

## Funding

J.A.B. was supported by the M. Guyer postdoctoral fellowship. E.V.W.S. was supported by a National Science Foundation Graduate Research Fellowship. C.E.S.L. was supported by postdoctoral CONACYT grant reg. 207146/454834. Morphological data collection was supported by Deutsche Forschungsgemeinschaft grant no. BR5039/3-1 to G.B. Fieldwork in Antarctica was supported by National Science Foundation grants ANT-0551969 to A.L.M. ANT-0440577 to H.A.W. Collection under projects POL2006-06399/CGL (CLIMANT) and CTM2012-39350-C02-01 (ECOWED) were funded by the Spanish Comisión Interministerial de Ciencia y Tecnología and the Ministerio de Economía y Competitividad. Revolta program 1124, led by Guillaume Lecointre, was supported by Institut polaire français Paul Émile Victor (IPEV). This work was supported by intramural funds of the American Museum of Natural History (W.C.W.) and by intramural funds of the University of Wisconsin-Madison and National Science Foundation grant no. IOS-1552610 (P.P.S).

## Acknowledgements

This work is dedicated to the memory of Roger Norman Bamber. We are indebted to Katsumi Miyazaki, Anna Soler-Membrives, and Julia D. Sigwart for providing specimens for study. We thank the crews of many research vessels for assistance with fieldwork, and the Australian and Paris Museums for loans of materials for study. Sequencing and assembly of the *Pycnogonum litorale* mitogenome was performed by Anke Braband and Jakob Machner. Chronograms of selected marine invertebrate taxa were kindly provided by Heather D. Bracken-Grissom, Simon Y.W. Ho, Tim D. O’Hara, Greg W. Rouse, and Jo M. Wolfe. Access to computing nodes for intensive tasks was facilitated the Bioinformatics Resource Center (BRC) and by Christina Koch at the Center for High Throughput Computing (CHTC) of the University of Wisconsin-Madison.

---

**Figure S2.** Comparative metrics of dataset performance by partition. Left column: Taxon sampling per locus. Middle column: Alignment length post-trimming. Right column: GC content.

**Figure S3.** Comparative metrics of dataset performance by partition. Left column: normalized Robinson-Foulds distance per locus (from species tree). Middle column: normalized weighted Robinson-Foulds distance per locus (from species tree). Right column: mean pairwise sequence identity (a proxy for evolutionary rate).

**Figure S4.** Maximum likelihood supermatrix analyses of individual data partitions. Colors in branches correspond to families (left). Annotated tree files with nodal support frequencies are available on the Dryad Digital Repository.

**Figure S5.** BAMM analysis of sea spiders showing lack of evidence for rate shifts (prior on expected number of rate shifts set to 10).

**Table S1.** Locality and accession data for study specimens.

**Table S2.** GenBank accession data for sequenced terminals (cells marked “XXX” are new sequence data under embargo).

**Text S1.** Extended materials and methods.

**Text S2.** Discussion of sea spider fossil record and implementation of phylogenomic dating.

**Text S3.** Comparative performance of phylogenetic data classes.

## Notes

#### Summary of Updates

A new supplementary figure has been added to show the results of phylogenetic inference using PhyloBayes-mpi. A different chronogram for Decapoda has been depicted in Figure 4b, on the counsel of the authors of reference 49. Two incorrect references have been corrected.

