## Supplementary Figures S1-S5 for "Phylogenomic resolution of sea spider diversification through integration of multiple data classes"

**Figure S1.** Tree topologies from PhyloBayes-mpi analyses of Matrices 1 and 2. Note the lack of convergence in both matrices (convergence is defined as *maxdiff* < 0.1 and minimum ESS > 200 for all summary statistics). Fully annotated tree topologies with nodal support values are available on the Dryad Digital Repository.

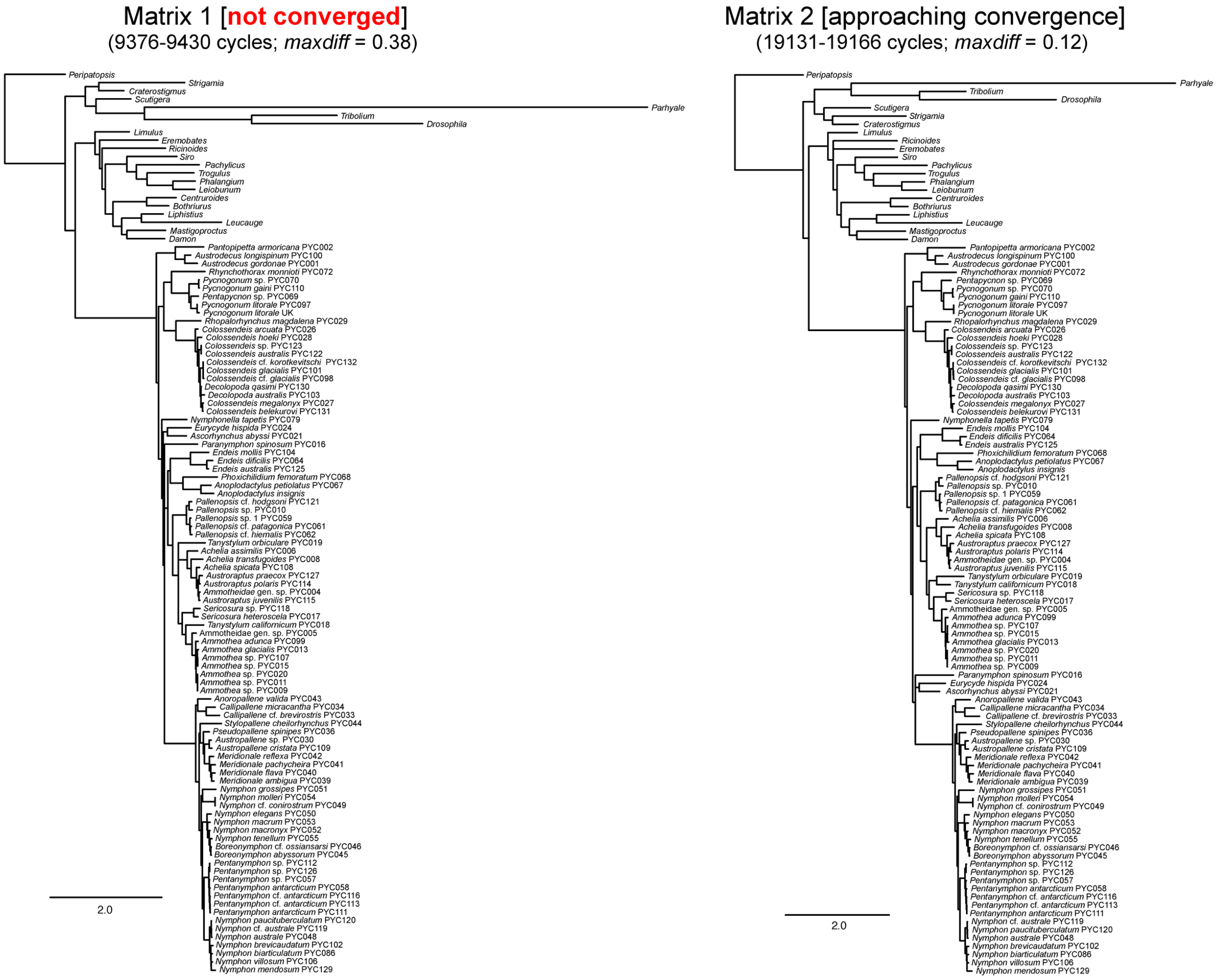

**Figure S2.** Comparative metrics of dataset performance by partition. Left column: Taxon sampling per locus. Middle column: Alignment length post-trimming. Right column: GC content.

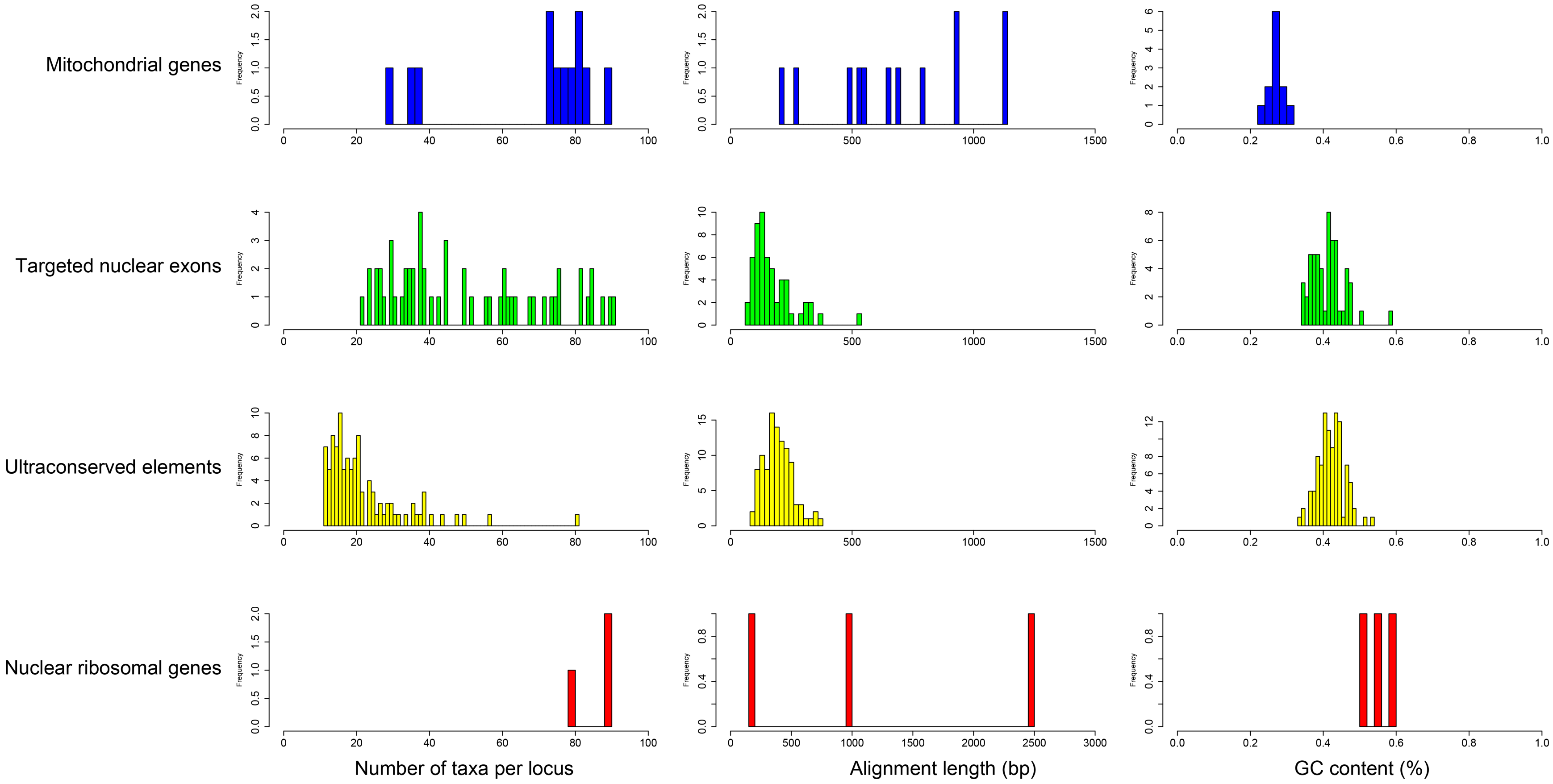

**Figure S3.** Comparative metrics of dataset performance by partition. Left column: normalized Robinson-Foulds distance per locus (from species tree). Middle column: normalized weighted Robinson-Foulds distance per locus (from species tree). Right column: mean pairwise sequence identity (a proxy for evolutionary rate).

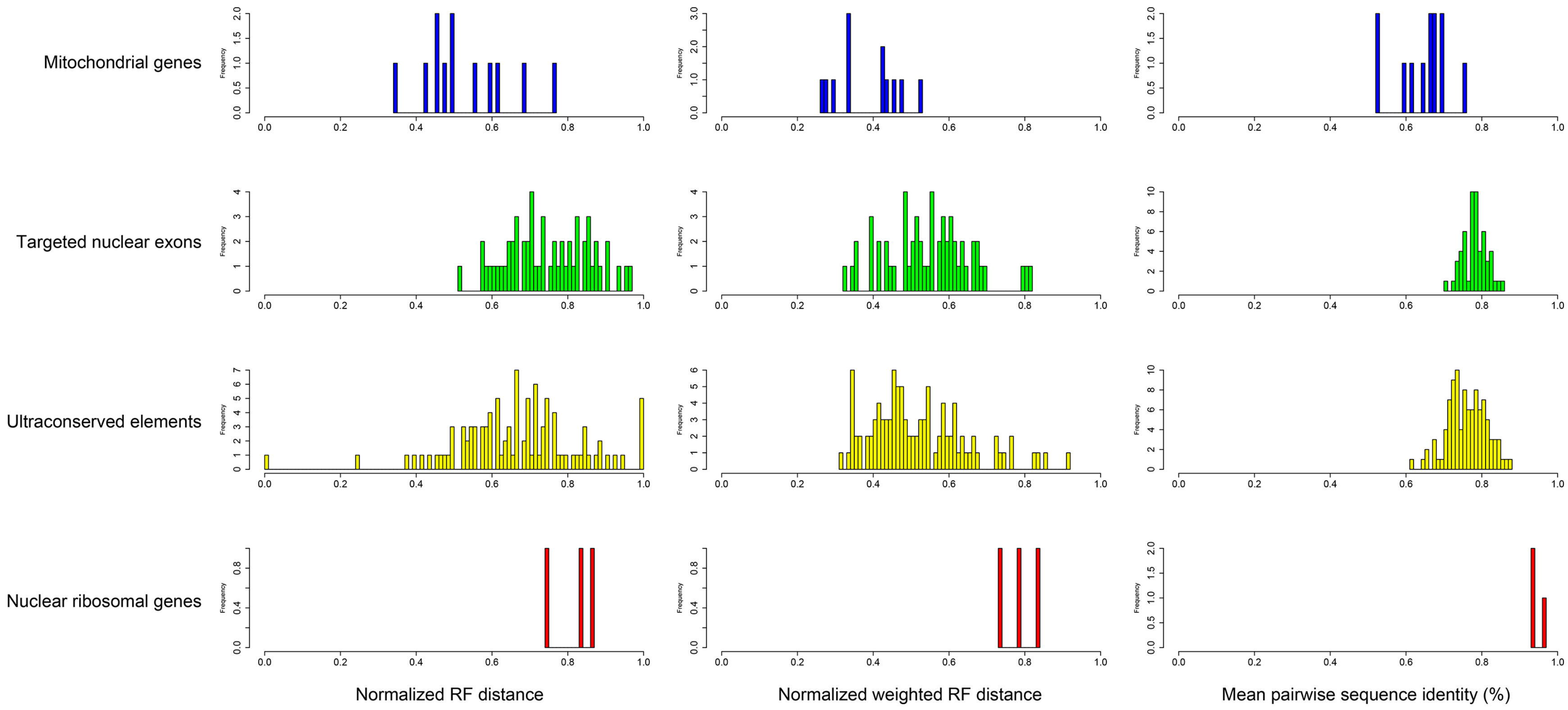

**Figure S4.** Maximum likelihood supermatrix analyses of individual data partitions. Colors in branches correspond to families (left). Annotated tree files with nodal support frequencies are available on the Dryad Digital Repository.

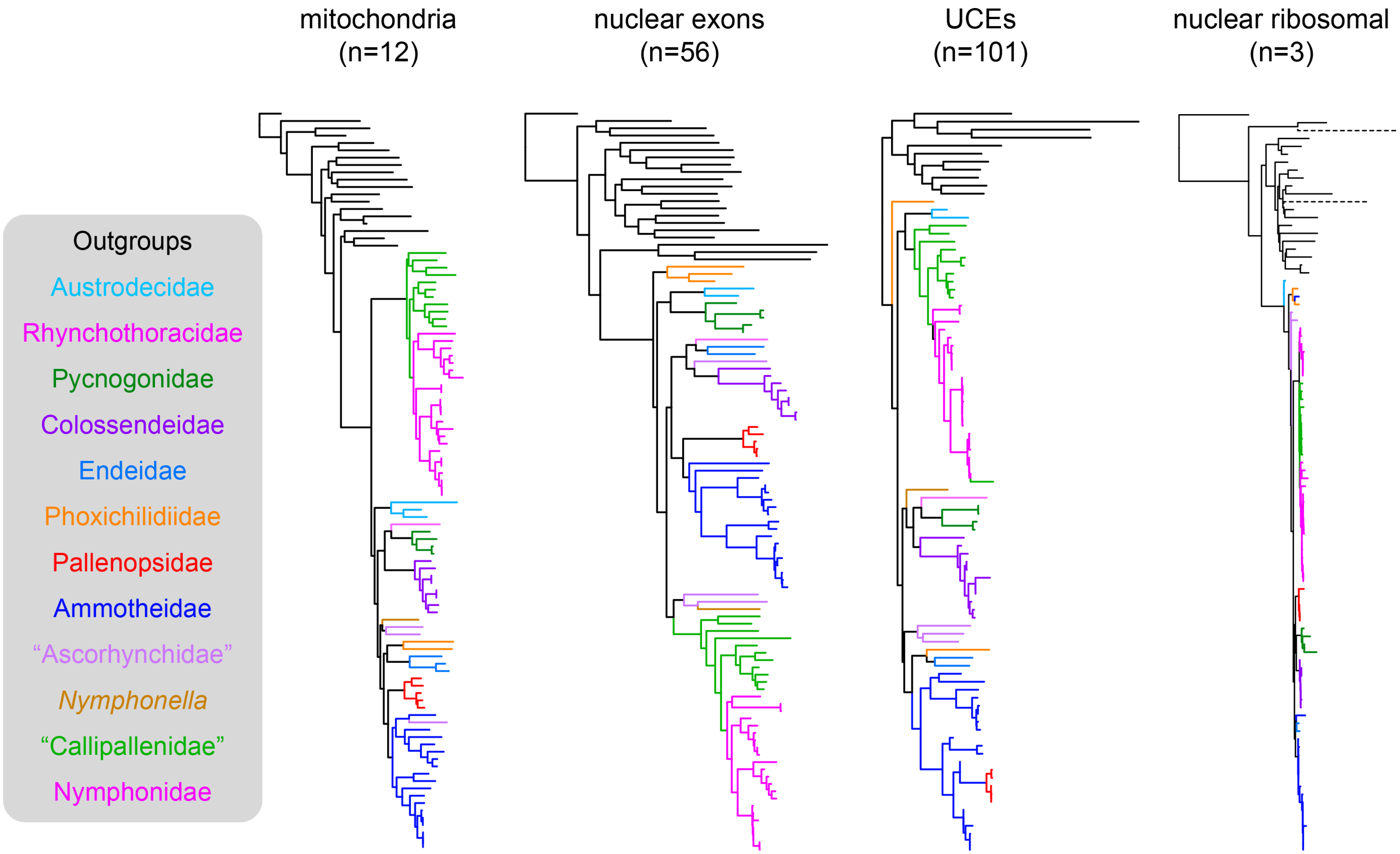

**Figure S5.** BAMM analysis of sea spiders showing lack of evidence for rate shifts. Outgroups removed prior to analysis.

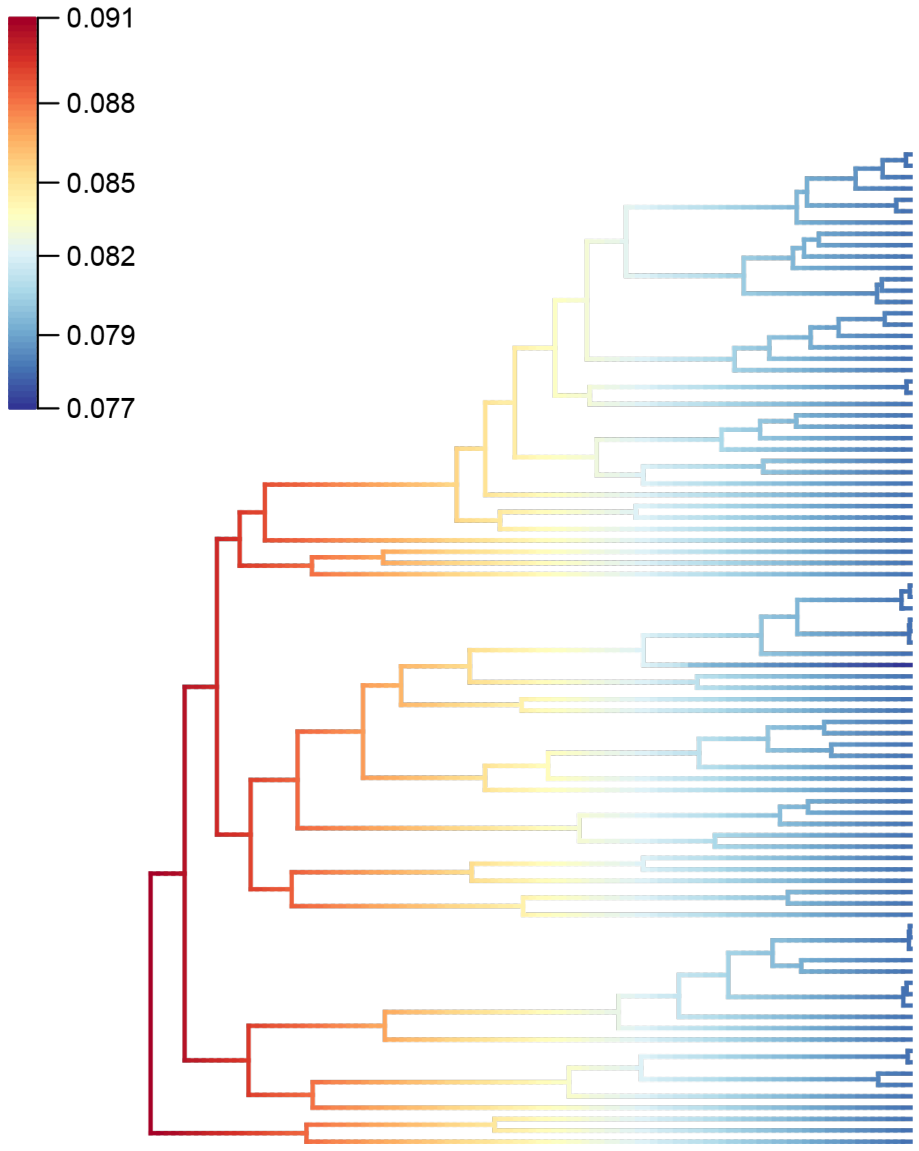
