## Supplementary Texts S1-S3 for "Phylogenomic resolution of sea spider diversification through integration of multiple data classes"

**Electronic supplementary material**

**Supplementary Text S1**: Extended materials and methods.

*Taxonomic sampling and probe design*

Specimens of all sea spider families were obtained from field expeditions and museum collections, as well as material collected during multiple deep sea cruises. A list of taxa sampled is provided as electronic supplementary material, table S1. DNA was extracted using the Qiagen DNeasy Blood and Tissue Kit (Valencia, CA, US) from one to five specimens, prioritizing tissue from the propodus and dactylus (gut-free leg podomeres) where possible. Taxonomic sampling consisted of 89 sea spiders; outgroups consisted of 14 Arachnida (including one Xiphosura, which has recently been shown to be nested within the arachnids [1, 2]), three Myriapoda, three Pancrustacea, and one Onychophora, as listed in electronic supplementary material, table S1.

Probes were designed using available references and databases, given the absence of a published sea spider genome. Probes for mitochondrial genomes were designed using an unpublished mitogenome of *Pycnogonum litorale* and five published sea spider mitogenomes: *Tanystylum orbiculare* (GU370074.1), *Ammothea hilgendorfi* (GU370075.1), *Achelia bituberculata* (NC_009724.1), *Nymphon unguiculatum-charcoti* complex sp. (GU370076.1), and *Nymphon gracile* (NC_008572.1). From previous work on the phylogeny of Chelicerata [3], we identified 56 nuclear exons with clock-like evolutionary rates that exhibited sufficient conservation for probe design with 3× tiling. Probe sequences are provided in the Dryad Digital Repository. For ultraconserved element (UCE) sequencing, we deployed probes from the UCE Arachnida 1.1Kv1 bait set [4]. Probe synthesis, automated library preparation, and paired-end sequencing (2 × 150 bp) was performed on the Illumina Hi-Seq 2500 platform through RAPiD Genomics (Gainesville, FL, US).

*Bioinformatic and phylogenomic operations*

Raw reads were assembled using Trinity v.2.7 [5] using a path reinforcement distance of 75. Reference sequences of sea spiders (mitogenomes and nuclear exons) and/or euchelicerates (UCEs) were used to separate loci into alignments. We included the nuclear ribosomal genes 5.8S rRNA, 18S rRNA, and 28S rRNA, which were obtained as bycatch; on-target amplification of these bycatch loci was verified using BLASTn against available sea spider ribosomal sequences in GenBank. Outgroup taxa were subsequently added into the alignments using available transcriptomes from our previous works (for nuclear exons [1-3]), from GenBank data (mitochondrial genes/genomes and nuclear ribosomal genes), and from a subset of the taxa in arachnid UCE alignments [4]. Due to the unavailability of the same phylogenetic data classes among the genomic resources of some data-poor outgroups, we created chimeric terminals for such terminals as *Eremobates* and *Ricinoides*. The nature of these congeneric chimeras is that sequence data for nuclear exons and ribosomal genes are drawn from the transcriptomes of one species (e.g., *Eremobates* cf. transfugoides; *Ricinoides atewa*), but the mitochondrial genome is drawn from a congener (e.g., *Eremobates* cf. *palpisetulosus*; *Ricinoides karschii*). The list of chimeric outgroup terminals is detailed in electronic supplementary material, table S2.

Multiple sequence alignment was performed using MUSCLE v.3.8.31 [6]; for nuclear and mitochondrial exons, alignments were performed using peptide translations and subsequently reverted to nucleotides. We discarded the mitochondrial genes 12S rRNA, 16S rRNA, and tRNA, due to their high rate of evolution, as well as off-target capture of nonsea spider sequence for these regions, which we presumed to have resulted from sequencing of gut content and/or epibionts.

Alignments were trimmed using trimal v.1.2 [7] to cull overhanging ends. Each alignment was filtered to ensure the inclusion of a minimum of six sea spider terminals and four outgroup terminals (resulting in the exclusion of the mitochondrial protein ATP8 and 129 out of 230 UCE alignments). Gene trees were inferred using IQ-TREE v.1.6 [8] with the best-fitting model selected by ModelFinder (-m MFP) [9]. Gene tree topologies were visually inspected for potential paralogs, which were summarily discarded from alignments (principally found in the UCE alignments previously generated for arachnids).

Four matrices were constructed for our main analyses. Matrices 1-3 were constructed using taxon occupancy thresholds of 50%, 33%, and 25%, respectively. Matrix 4 consisted of loci stipulated to sample at least one Austrodecidae (the putative sister group to the remaining sea spiders). The final alignments were analyzed as concatenated supermatrices using IQ-TREE v.1.6 with best-fitting models per locus. Due to ongoing debate over the benefits of model-fitting, we additionally analyzed our datasets using a unique GTR + $\Gamma_4$ model for each locus as well. Nodal support was estimated using bootstrap resampling frequency with 1000 ultrafast bootstrap replicates in IQ-TREE. In addition, Approximately Unbiased (AU) tests of monophyly were performed using in-built tools in IQ-TREE for selected phylogenetic hypotheses using Matrices 1 and 4.

For assessment of dataset performance, we inferred the normalized Robinson-Foulds (RF) and weighted Robinson-Foulds (wRF) distances between the species tree inferred using the most complete matrix (Matrix 1) and each gene tree. To overcome missing data, the species tree was iteratively pruned of tips to match the terminals of each gene tree prior to calculation of RF and wRF distances, and the distances were normalized using the maximal possible distance between each gene tree and the pruned species. Other metrics of performance consisted of locus length, taxon sampling, GC content, and mean pairwise sequence identity (MPSI; a proxy for evolutionary rate). Tree manipulations and calculations of performance metrics were performed using R packages ape v.5.3 [10] and phytools v.0.6-99 [11].

*Bayesian inference analysis*

PhyloBayes-mpi v.1.7 [12] analysis was performed under the CAT + GTR + $\Gamma_4$ model for Matrices 1 and 2 (the two most complete matrices). For each matrix, two runs were performed lasting 9376 to 19166 cycles across all analyses. Convergence was assessed using inbuilt tools in PhyloBayes-mpi v.1.7, as well in Tracer v.1.7 [13]. As a conservative measure, the intial 10% of all runs were discarded as burnin. Tree topologies and convergence assessment metrics are reported in electronic supplementary material, figure S1.

*Phylogenomic dating*

Phylogenomic estimation of divergence times was estimated using a node dating approach with MCMCTree [14] on two datasets (Matrices 1 and 3), implementing a likelihood approximation of branch lengths using a multivariate normal distribution [15]. Fossils used to inform the dating consisted of 11 outgroup and four ingroup node calibrations (the Silurian sea spider *Haliestes dasos* and four Jurassic fossils representing the families Endeidae, Colossendeidae, and Ammotheidae). All calibrations were implemented as soft minimum and soft maximum ages. A detailed justification of these calibrations and their use (as well as an overview of fossils excluded for this purpose) is discussed in electronic supplementary material, text S2.

Both the independent rates and correlated rates clock models were used to infer node ages with Matrices 1 and 3, under the maximum likelihood tree topology inferred for Matrix 3 with a unique substitution model per locus (as shown in figure 3). Four independent chains were run for $10^6$ generations for each analysis, sampling every $100^{th}$ generation. Convergence diagnostics were assessed using Tracer v.1.7 [13] and inbuilt tools in MCMCTree, ensuring minimum effective sample sizes >200 for summary statistics. As a conservative measure, 1000 trees (10%) were discarded as burnin.

Analyses of diversification rates through time were performed using Bayesian Analysis of Macroevolutionary Mixtures (BAMM v.2.5.0) [16]. Outgroups were removed prior to analysis using phytools. Rate shifts were permitted on all branches. MCMC chains were run for 500 000 generations, sampling event data every $10^4$ generations, and discarding 25% of the run as burnin.

*Micro-computed tomography*

Fixed specimens were dehydrated via an ascending ethanol series, incubated in solution of 2% iodine (resublimated; Carl RothGmbH&Co. KG, Karlsruhe, Germany; cat. #X864.1) in 99.5% ethanol for ca. 48 hours at room temperature, briefly rinsed in 99.5% ethanol and either transferred into a vial in ethanol (“wet scan”) or critical point-dried with a Leica EM CPD300 and glued on plastic welding rods (“dry scan”). Scans were performed with an

Xradia MicroXCT-200 (Carl Zeiss Microscopy GmbH). Settings were individually optimized for each specimen, including objective choice (0.39x, 4x, 10x) according to size. Scans were performed under 40kV/200µA/8W, with Binning 2 (noise reduction) and exposure times ranging from 0.4-1.5 sec. Tomography projections were reconstructed using the XMReconstructor software (Carl Zeiss Microscopy GmbH) with Binning 1 (full resolution) and TIFF format image stacks as output. Processing and visualization of image stacks (including highlighting of cephalic appendages and removal of non-target structures) was performed with Imaris (version 7.0.0., Bitplane AG, Switzerland) as described previously [17].

**Supplementary Text S2**: Discussion of sea spider fossil record and implementation of phylogenomic dating.

*The sea spider fossil record*

The oldest unequivocal fossil of crown group Pycnogonida, *Haliestes dasos*, dates to the Silurian (424 Mya) [1], whereas the oldest fossil assigned to the pycnogonid (stem) lineage is an early developmental instar from the Upper Cambrian (*Cambropycnogon klausmuelleri*; 501 Mya) [2]. Owing to the completeness of this fossil, the quality of preservation, and the detail of its three-dimensional scan, it is evident that the chelifores of *H. dasos* resembles those of Nymphonidae. The lack of an annulated or segmented proboscis rule out a placement close to the families Austrodecidae and lineages like *Eurycyde*. Moreover, *H. dasos* shares with extant sea spiders the reduced, unsegmented condition of the vestigial opisthosoma. Given that this fossil has been recovered at the base of Pycnogonida in some morphological cladistic analyses [1], we calibrated the age of crown group Pycnogonida using a soft minimum age of 424 Mya and a soft maximum age of 501 Mya.

*H. dasos*, however, is not considered the oldest sea spider fossil. The fossil *Palaeomarachne granulata* is Ordovician in age and is putatively a sea spider as well [3]. *P. granulata* is an odd specimen that exhibits a segmented region anterior of the cephalon, a condition unknown in any other sea spider (including fossil groups). In the present study, we excluded the use of this fossil from node dating owing to its poor preservation and incompleteness. Moreover, we are not convinced that *P. granulata* is a sea spider, or even a chelicerate arthropod, as no definitive characters specific to Chelicerata are visible in this fossil.

Two well-preserved Devonian fossils that clearly comprise sea spider stem-groups are *Flagellopantopus blocki* [4] and *Palaeoisopus problematicus* [5], as both exhibit a segmented opisthosoma and a well-developed telson—features considered plesiomorphic for Chelicerata but lost in extant Pycnogonida (figure 4*a*). While its cephalon is not well preserved, *F. blocki* exhibits an elongate flagellum with numerous articles, a condition observed in extant chelicerate groups like Palpigradi and Thelyphonida, as well as the recently described Cretaceous uraraneid, *Chimerarachne yingi* [6]. *F. blocki* and *P. problematicus* are suggestive of greater body plan diversity in Paleozoic stem lineages of Pycnogonida, but do not contribute greatly to the implementation of node dating, as *H. dasos* is older than both these fossils. For this reason, we did not use these Devonian taxa for node calibration.

Other Devonian sea spiders (*Palaeothea devonica*, *Palaeoisopus devonicus*, *Palaeopantopus maucheri*, and the ten-legged *Pentapantopus vogteli*) [5,7] were not considered usable for node dating, due to ages younger than *Haliestes dasos*, and in some cases, poor preservation of morphology.

Sea spider crown group fossils from the Late Jurassic of Solnhofen (160 Mya) were used to calibrate ingroup nodes: *Palaeopycnogonides gracilis* (Ammotheidae), *Colossopantopodus boissinensis* and *Colossopantopodus nanus* (Colossendeidae), and *Palaeoendeis elmii* (Endeidae) [8,9]. Each of these fossils was treated as a minimum age calibration for the stem-age of the corresponding family, with a maximum soft bound age of 501 Mya. The choice of constraining stem-age versus crown-age of the families was determined through preliminary runs assessing the fit of the prior distributions, after running MCMCTree with no data.

?*Eurycyde golem* was not used in this study, as the poor preservation precludes reliable placement in any extant sea spider genus or family [9].

*Outgroup calibrations*

For Arachnida, we constrained the crown group of Opiliones using the soft minimum age of 411 Mya, based on the harvestman fossil *Eophalangium sheari*, and the divergence of Eupnoi harvestmen with a soft minimum age of 305 Mya, based on the age of *Ameticos scolos* [10]. The divergence of spiders from the remaining Arachnopulmonata was constrained to at least 386 Mya (age of *Attercopus fimbriunguis*) and the divergence of *Liphistius* at 305 Mya (age of *Palaeothele montceauensis*) [6,11]. Pedipalpi (Amblypygi + Uropygi) was set to a minimum age of 319 Mya, based on the age of the fossil *Parageralinura naufraga* [12]. The divergence of Xiphosura from Riniculei was constrained to a minimum of 445 Mya (age of *Lunataspis aurora*) [13]. The stem-group age of scorpions was constrained to a soft minimum age of 435 Mya and a soft maximum of 514 Mya (based on the ages of *Parioscorpio venator* and *Eramoscorpius brucensis*) [14,15]. The crown group age of scorpions was constrained between 112.6 and 313.7 Mya (based on the ages of *Protoischnurus axelrodurum* and *Compsoscorpius buthiformis*) [16].

For Mandibulata, we constrained the basal diversification of Chilopoda between 416 and 521 Mya, and of Pleurostigmophora between 382.7 and 521 Mya. Altocrustacea (the node comprising the most recent common ancestor of the malacostracan exemplar *Parhyale hawaiensis* and the hexapods in this analysis) was constrained between 429.8 and 521 Mya. The root of the tree (split of Onychophora and Arthropoda) was constrained between 550 and 636 Mya. Mandibulate and root calibrations are based on various stem-group arthropod and lobopodian fossils reviewed by Wolfe et al. [17].

**Supplementary Text S3**: Comparative performance of phylogenetic data classes.

Performance measures for phylogenetic data classes consisted of number of taxa sampled, alignment length, GC content, RF distance, weighted RF distance, and evolutionary rate. The mean number of taxa per locus was highest for the mitochondrial and nuclear ribosomal genes (68.2 and 85.7, respectively) and comparable for targeted exons, albeit with high variance (mean=50.9, $\sigma^2$=20.9). UCE loci bore the most missing data, with an average of 22.2 terminals per locus. In addition, after trimming, the targeted exons and UCEs had comparable alignment lengths, and were shorter on average than mitochondrial and nuclear ribosomal genes. GC content of nuclear exons and UCEs was also comparable, with the average values intermediate between mitochondrial genes and nuclear ribosomal genes (electronic supplementary material, figure S2).

For assessment of dataset performance, we inferred the normalized Robinson-Foulds (RF) and weighted Robinson-Foulds (wRF) distances between the species tree inferred using the most complete matrix (Matrix 1) and each gene tree. To overcome missing data, the species tree was iteratively pruned of tips to match the terminals of each gene tree prior to calculation of RF and wRF distances, and the distances were normalized using the maximal possible distance between each gene tree and the pruned species. Assessment of congruence between gene trees and the pruned species trees showed that targeted exons and UCEs had comparable distributions of RF and wRF distances (electronic supplementary material, figure S3). Average values of both metrics were intermediate for targeted exons and UCEs, with respect to the lower values of mitochondrial genes and the higher values of nuclear ribosomal genes. Paralleling these distributions, MPSI values of targeted exons and UCEs were intermediate between mitochondrial genes and nuclear ribosomal genes (electronic supplementary material, figure S3).

Inference of tree topologies based on each of the four data partitions was performed in IQ-TREE with identical heuristics as for the species tree analyses. Comparison of tree topologies reflected the performance metrics measured above; mitochondrial genes and targeted exons retained many of the higher-level relationships recovered by the species tree, such as the monophyly of various families, and the sister group relationships of Callipallenidae + Nymphonidae and Pallenopsidae + Ammotheidae (electronic supplementary material, figure S4). However, by themselves, none of these partitions was able to resolve basal relationships with support. Reflecting the high proportion of missing data, the UCE tree topology was largely discordant with the species tree and other partitions, although some higher-level relationships were consistently recovered (e.g., the monophyly of some families; Callipallenidae + Nymphonidae; Pallenopsidae + Ammotheidae). The nuclear ribosomal partition retained some higher-level relationships, but branch lengths subtending interfamilial relationships were close to zero and were not supported (electronic supplementary material, figure S4).
